# Sex Differences in Cachexia Outcomes and Branched-Chain Amino Acid Metabolism Following Chemotherapy in Aged Mice

**DOI:** 10.1101/2025.10.30.685610

**Authors:** Stephen Mora, Gagandeep Mann, Olasunkanmi A J Adegoke

## Abstract

Cachexia is a complex muscle wasting syndrome that affects majority of hospitalized cancer patients receiving chemotherapy. It is often unresponsive to nutritional interventions, including provision of branched-chain amino acids (BCAA: leucine, isoleucine and valine). BCAA are anabolic for skeletal muscle. We wondered whether their ineffectiveness in managing cachexia might be related to altered metabolism of these amino acids, a subject that has received minimal attention. Because estrogen limits BCAA catabolism, we hypothesized that the effects of chemotherapy on cachexia in old mice would be worse in males compared to females, and that this would be related to greater tissue release of BCAA in males. To better reflect age population for which cachexia is an issue, we treated aged male and female mice (18±2 months) with the chemotherapy drug cocktail FOLFIRI (50mg/kg 5-fluorouracil (5FU), 90mg/kg Leucovorin, and 24mg/kg CPT11) or vehicle twice per week for 6 weeks. This cocktail is used in treating colon cancer. Metabolism and concentrations of the BCAA and their metabolites were measured in plasma and tissues. There was a main effect of chemotherapy, reflected in reduced body weight, skeletal muscle, myofibrillar protein content, anabolic signalling and protein synthesis. In response to chemotherapy, males showed worsened outcomes for skeletal muscle weight and ubiquitinated proteins; they also had higher total plasma BCAA but reduced muscle BCAA. There was a main effect of chemotherapy in reducing the expression of the BCAA transporter LAT1. In response to chemotherapy, gastrocnemius muscle of males but not females had reduced inhibitory phosphorylation of BCKD-E1α^ser293^, corresponding with increased activity of this enzyme. Chemotherapy reduced muscle and liver ketoacids of the BCAA only in females. These data suggest that sex differences in BCAA catabolism may be linked to the severity of chemotherapy-induced muscle damage and interventions against cachexia need to take this into account.

## Introduction

Chemotherapy drugs are a main line of treatment for diverse cancers. However, their use is associated with alterations and damage to diverse body organs and tissues leading to nausea and vomiting, reduced appetite, neutropenia, loss of hair, symptoms related to cardiotoxicity, and neurotoxicity, and diverse others (1–4). FOLFIRI (a mix of **FOL**inic acid, 5-**F**luorouracil (5FU) and **IR**Inotecan (CPT11)) is a chemotherapy drug cocktail that is used in treating one of the most common cancers, colorectal cancers (5–7). In addition to the side effects mentioned above, data from *in vitro* and rodent studies show specific detrimental effects of the cocktail on skeletal muscle and myotubes, including reduced muscle mass, fiber cross-sectional area and strength (8–10), reduced muscle protein synthesis (10), increased expression and activities of markers of proteolysis (10), reduced abundance and function of the mitochondria (11), reduced myotube diameter and abundance of myofibrillar proteins (12,13), altered substrate metabolism (8), and increased reactive oxygen species (8,14). In clinical studies, use of chemotherapy regimens that include 5FU or CPT11, which are components of FOLFIRI, is associated with skeletal muscle loss (15). Also, cases of muscle twitching have also been reported in colorectal patients on FOLFIRI (16–18).

Damage and toxicity to the muscle can contribute to cachexia, a complex metabolic body and muscle wasting syndrome that affects the majority of hospitalized cancer patients receiving chemotherapy (19,20). This wasting syndrome is associated with poor treatment outcomes, treatment-related toxicities, poor quality of life, and mortality (21). In skeletal muscle, cachexia is associated with impaired activation of the mammalian/mechanistic target of rapamycin complex 1 (mTORC1), a complex that is anabolic for skeletal muscle (mTORC1) (22,23). Cachexia is also associated with activation of muscle catabolic pathways, including components of ubiquitin-proteasome and autophagy pathways (24–27). A main challenge of cachexia is that there is currently no effective therapy for the condition. Unlike starvation where provision of nutrients will suffice to correct the condition, cachexia is often refractory to nutritional interventions. This may relate to impaired nutrient metabolism, especially the metabolism of amino acids which are the macronutrients that are vital to body anabolism.

Branched-chain amino acids (BCAA: leucine, isoleucine and valine) are transported into the skeletal muscle by the L-type amino acid transporter 1 (LAT1). During their catabolism, BCAA are transaminated by branched-chain aminotransferase 2 (BCAT2) into their respective branched-chain α-ketoacids (BCKA): 2-keto-isocaproate(4-methyl-2-oxopentanoic acid (KIC)) from leucine, α-keto-β-methylvaleric acid (3-methyl-2-oxopentanoate (KMV)) from isoleucine, and 2-keto-isovalerate(3-methyl-2-oxobutanoic acid (KIV)) from valine. BCKA are then oxidatively decarboxylated by branched-chain α-keto acid dehydrogenase complex (BCKD) (28), generating metabolites such as acetyl-CoA and succinyl-CoA that can be funneled into diverse metabolic pathways. In addition, leucine-derived KIC can be metabolized to beta-hydroxymethylbutyrate in the liver, a metabolite that is anabolic for skeletal muscle (29,30).

Some animal studies have shown that BCAA have positive effects on reversing cancer-induced cachexia, but not survival (31–33). BCAA also offer minimal effectiveness in treating cachexia in humans (34–37), which may be related to limitations of pre-clinical models and little understanding of amino acid metabolism in cachexia. *In-vivo* rodent models that study cancer-and/or chemotherapy-induced cachexia investigate mainly young male mice ((38–40), S1 Table). Thus, not much is known about BCAA metabolism in old animals. Also, sex differences exist for BCAA metabolism. For example, compared to female individuals, males have higher skeletal muscle BCAA concentrations (41) and greater leucine oxidation following endurance exercise (42). In addition, estrogen reduces BCAA catabolism in female rats (43). Aged individuals (44,45) and animals (46) exhibit lower circulating amino acid concentrations and reduced dietary protein intake compared to their younger counterparts. However, the effect of chemotherapy on the abundance and the activity of enzymes involved in BCAA catabolism have not been investigated in any aged animal tissues. This is an important question because the diminished effectiveness of BCAA in attenuating skeletal muscle loss in cachexia may be related to chemotherapy-induced alterations to skeletal muscle amino acid metabolism. Because estrogen limits BCAA catabolism (43), we hypothesized that the effects of chemotherapy on indices of cachexia in old mice would be worse in male compared to female mice, and that this would be related to greater tissue release of BCAA (through protein breakdown) in male compared to female animals. We present data on sex differences in cachexia outcomes and BCAA metabolism following chemotherapy in aged mice, as chemotherapy treatments are a cause of cachexia in animals (38–40) and clinically (47–49).

## Materials and methods

### Mice

All animal experiments were approved by the York University Animal Care Committee (Protocol #2020-09) and were conducted in line with the guidelines of the Canadian Council on Animal Care. Twenty male and 20 female 8-week-old CD2F1 mice were purchased from Charles River Laboratories. Mice were acclimatized and housed in the vivarium with free access to food (Purina 5015*, LabDiet, St. Louis, MO) and water. They were aged until 18±2 months prior to treatment. Male and female mice were administered intraperitoneally either the chemotherapy drug cocktail FOLFIRI (50mg/kg 5FU (#F6627), 90mg/kg Leucovorin (#F7878) and 24mg/kg CPT11 (#I1406); drug) or vehicle (3.8% DMSO (#D5879) in saline), all from Sigma Aldrich, St. Louis, MO, twice per week for 6 weeks (n=10 for all groups). Weekly dosages were separated by two days. This chemotherapy drug cocktail is used to treat colorectal cancer (50). The doses used were based on a previous study (39) and deemed to not exceed clinically relevant concentrations (38). Body weight and food intake were recorded daily. Body weight was expressed as weekly variations relative to baseline as presented previously (38). Twenty-four hours (h) after their last chemotherapy dose, mice were starved for 3 h prior to being euthanized via cervical dislocation. Several skeletal muscles (gastrocnemius, tibialis anterior (TA), and quadriceps) and other tissues (e.g. heart, kidney, liver, spleen and visceral adipose tissue) were collected, weighed, flash-frozen in liquid nitrogen and stored at −80°C until analysis. Animal handling and tissue harvesting were conducted in a careful manner to minimize stress and pain as instructed by institutional vivarium staff.

### Insulin Tolerance Test

On the days when insulin tolerance test (ITT) was done, at least 24 h from their previous chemotherapy dose, mice were starved for 6 h. Blood samples were collected from the saphenous vein on glucose strips (Alpha TRAK, #71681, Parsippany, NJ) and inserted into a glucometer (Alpha TRAK, #71675-01) at 0 (baseline) and at 5, 15, 30 and 120 minutes after a subcutaneous insulin injection (0.75units/kg; Eli Lilly, Humulin R, #00586714, Indianapolis, IN). ITT was performed at weeks 3 and 6 of chemotherapy treatment.

### Protein Synthesis (SUnSET Analysis)

With at least 24 h from their last chemotherapy dose, as previously described (51), mice were starved for 3 h and then intraperitoneally injected with 0.040μmol/g bodyweight of puromycin (Sigma Aldrich, #P8833) in saline 30 minutes prior to euthanasia. After euthanasia, skeletal muscle (gastrocnemius) proteins were immunoblotted against an anti-puromycin antibody and corrected to their Ponceau S staining.

### Western Blotting

Gastrocnemius muscle and liver were homogenized in 7X complete buffer: 20mM HEPES, 2mM EGTA, 50mM NaF, 100mM KCI, 0.2mM EDTA, 50mM B-Glycerophosphate. This buffer was supplemented with protease inhibitor (10µL/mL; Sigma Aldrich, #P8340), phosphatase inhibitor (10µL/mL; Sigma Aldrich, #P5726), 0.2M sodium vanadate (2.5µL/mL), 1M DTT (1µL/mL) and 0.2M benzamidine (5µL/mL) prior to use. Homogenates were then centrifuged (1000g for 3 minutes at 4°C), followed by the removal of the supernatant and further centrifugation (10000g for 30 minutes at 4°C). The Pierce BCA Protein Assay kit (Thermo Scientific, #23225, Waltham, MA) was used to determine protein concentrations in the resulting supernatant. Equal amounts of protein were separated on 10 or 15% SDS-PAGE gels and transferred onto polyvinylidene difluoride membranes (0.2μM, BIO-RAD). Incubation of membranes in primary (S2 Table) and secondary antibodies (HRP-conjugated anti-rabbit (#7074) or anti-mouse (#7076), Cell Signalling Technology, Danvers, MA), imaging and quantification of data were as described (52–54).

### Amino Acid and Keto Acid Concentrations

Plasma, gastrocnemius muscle and liver amino acid levels were measured as previously described (55). Tissues were homogenized (Bio-Gen PRO200 Homogenizer, PRO Scientific, Oxford, CT), centrifuged and the supernatant (containing all the amino acids) was diluted and pre-column derivatized in a ratio of 1 (sample): 1 (o-phthalaldehyde, Sigma Aldrich, #P1378). Each sample was then injected into a YMC-Triart C18 column fitted onto an ultra HPLC system (Nexera X2, Shimadzu, Kyoto, Japan) connected to a fluorescence detector (Shimadzu, Kyoto, Japan; excitation: 367 nm; emission: 446 nm). Dilution, pre-column derivatization and injection occurred similarly for plasma samples. Amino acid concentrations were calculated by generating amino acid standard (Sigma Aldrich, #AAS18) curves and by normalizing muscle and livers samples to total protein.

Ketoacids were measured as described by Fujiwara et al (56). Gastrocnemius and liver samples were homogenized as above. Plasma samples and tissue homogenates were then diluted and treated with 1,2-diamino-4,5-methylenedioxybenzene (DMB, Sigma Aldrich, #66807) in a 1:1 ratio. A 1mL solution of DMB was made by mixing 1.6mg DMB, 4.9mg sodium sulfite, 70μL of 2-mercaptethanol, and 58μL of concentrated HCl in 870μL of water. Samples were then heated (85°C, 45 minutes) before being cooled (on ice) and injected into a Inertsil ODS-4 column (2 μm, 100 × 2.1 mm; GL Sciences, Torrance, CA, USA) fitted onto an ultra high pressure liquid chromatography (UHPLC) system connected to a fluorescence detector. To elute the BCKA, mobile phases A (30% water, 70% MeOH) and B (100% MeOH) with a flow rate maintained at 0.2mL/minute and stable column temperature (40°C) were used. Values from skeletal muscle and liver samples were normalized to total protein concentrations.

### BCKD Activity Assay

We modified a previously described assay (57). Frozen gastrocnemius muscle and liver samples were homogenized in 250µL of ice-cold buffer 1 (30mM potassium phosphate buffer (KPI), 3mM EDTA, 5mM DTT, 1mM valine, 3% FBS, 5% Triton X-100, 1µm leupeptin (Sigma Aldrich, #L2884)). Following sample centrifugation (10 minutes at 10,000g, 4°C), the supernatant (50μL) was added into 300μL of buffer 2 (50mM HEPES, 30mM KPI, 0.4mM CoA (Sigma Aldrich, #C4282), 3mM NAD+ (Sigma Aldrich, #N0632), 5% FBS, 2mM Thiamine (Sigma Aldrich, #T1270), 2mM magnesium chloride and 7.8µM [^14^C]valine (Perkin Elmer, #NEC291EU050UC, Waltham, MA)). All reactions took place in a 1.5mL Eppendorf tube that had a raised wick trap that had been impregnated with 2M NaOH. The tube was then tightly sealed with tape and capped, prior to being placed in a shaking incubator (37°C for 30 minutes). The radiolabeled ^14^CO_2_ captured in the wick trap during the incubation was counted in a liquid scintillation counter. To calculate BCKD activity, we divided the counts per minute on each wick by the amount of counts that is equal to 1μmol of BCKD enzyme activity and corrected the resulting values for total protein.

### Statistical Analysis

All immunoblot data were adjusted to their corresponding γ-tubulin values. Graphs were drawn using Prism 10 (GraphPad software). A two-way analysis of variance (ANOVA) with a Tukey’s post-hoc test was used to measure the main effects of chemotherapy, the main effects of sex, and of the interactions between the two. Results were expressed as mean ± standard error of the mean (SE). Significance was determined as p-value < 0.05.

## Results

### Both Sexes Exhibit Body and Adipose Tissue Loss following Chemotherapy, but Muscle Wasting was More Severe in Males

There was a main effect of chemotherapy drugs with the drugs reducing body weight in both sexes (Fig 1A). For weekly mean body weights, there was a main effect of chemotherapy from week 4, which was more profound in weeks 5 and 6 (Table 1) as both sexes had reduced body weights. There was a main effect of sex on body weights during each of the six weeks studied, as males regardless of treatment group had greater body weight than females. In week three, control males had reduced body weight compared to chemotherapy-treated males. (Table 1).

**Fig 1:**
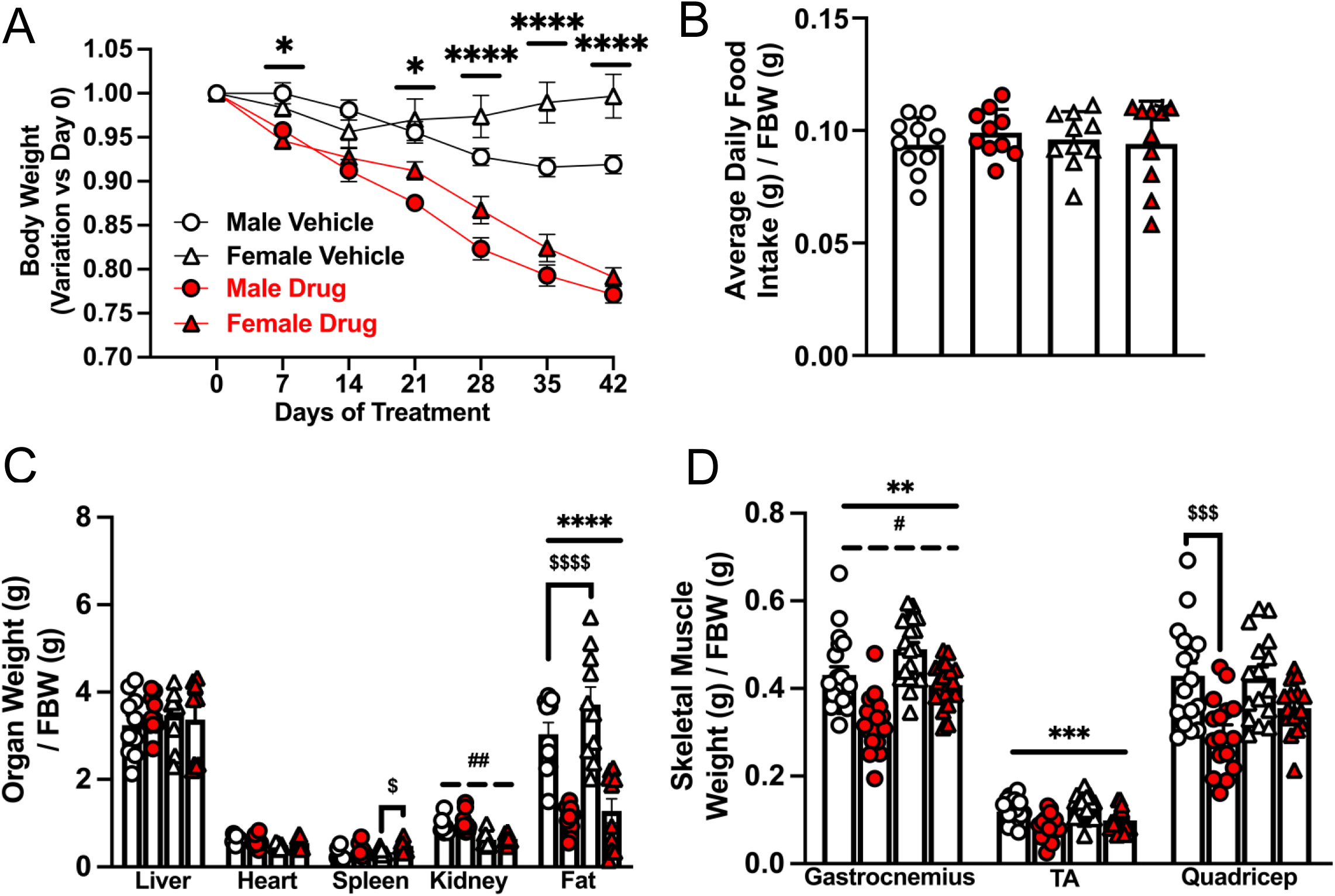
Greater Muscle Wasting in Males Compared to Females Following Chemotherapy Treatment in Old Mice. Male and female (18±2 months of age) CD2F1 mice were treated with either vehicle (males: white circle, females: white triangle; 3.8% DMSO in saline) or a chemotherapy drug cocktail (males: red circle, females: red triangle; 50mg/kg 5FU, 90mg/kg Leucovorin, 24mg/kg CPT11) twice per week for 6 weeks. Body weight relative to initial weight (A) and average daily food intake (B) were recorded for 6 weeks. At least 24 h after their last chemotherapy dose, mice were euthanized. Food intake, as well as the weights of the organs (C) and skeletal muscles (D) were normalized to FBW. Values for fat are for visceral adipose tissue. Data are mean ± SE, n = 8 – 10. Main effect of chemotherapy: ** p < 0.01, *** p < 0.001, **** p < 0.0001; main effect of sex: # p < 0.05, ## p < 0.01; interaction effect of chemotherapy and sex: $ p < 0.05, $$$ p < 0.001, $$$$ p < 0.0001. Data were analyzed using a two-way ANOVA followed by a Tukey’s post hoc test. In Fig 1A, two-way ANOVA analyses (chemotherapy, sex) were done at each time point. h, hours; FBW, final body weight; g, gram.

**Table 1.** Mean weekly body weight (g)

|  | Control |  | Drug |  | Statistics |  |  |
| --- | --- | --- | --- | --- | --- | --- | --- |
| Week | Male | Female | Male | Female | Treatment | Sex | Interaction |
| 1 | 45.1±0.9 | 37.2±1.6 | 42.6±0.9 | 34.5±0.9 |  | #### |  |
| 2 | 44.3±1.1 | 33.2±1.3 | 40.5±0.7 | 34.0±0.7 |  | #### |  |
| 3 | 43.2±1.1.0 | 33.5±1.2 | 37.7±1.6 | 33.5±0.8 | | ### | \$ |
| 4 | 41.9±1.0 | 33.8±1.2 | 36.4±0.7 | 28.1±0.8 | ** | #### |  |
| 5 | 41.4±1.0 | 37.2±1.2 | 35.1±0.7 | 26.6±0.8 | *** | #### |  |
| 6 | 41.5±1.0 | 37.4±1.3 | 33.5±0.5 | 23.4±0.6 | **** | #### |  |

Main effect of chemotherapy: \*\* p < 0.01, \*\*\* p < 0.001, \*\*\*\* p < 0.0001; main effect of sex: ### p < 0.001, #### p < 0.0001; interaction effects of chemotherapy and sex: \$ p < 0.05: in week 3, compared to control, chemotherapy reduced body weight only in male mice.

Chemotherapy-induced body weight loss was not due to differences in food intake (Fig 1B). For weekly mean food intake, there was no statistically significant main effect of chemotherapy or sex (Table 2). However, in weeks 2 and 3, chemotherapy-treated females ate more than control female mice. This was not observed in males. In week 5, chemotherapy-treated female mice ate less than control female mice and less than chemotherapy-treated male mice (Table 2). Chemotherapy had no effect on measured daily food intake (S1Table), consistent with data in Fig 1B and Table 2. On some days, male mice ate more than female ones. As the study progressed, there was an interaction effect in that chemotherapy-treated females ate less than control females.

**Table 2.** Mean weekly food intake per body weight (mg/g)

|  | Control |  | Drug |  | Statistics |  |  |
| --- | --- | --- | --- | --- | --- | --- | --- |
| Week | Male | Female | Male | Female | Treatment | Sex | Interaction |
| 1 | 65±7 | 78±7 | 61±4 | 74±3 |  |  |  |
| 2 | 107±8 | 109±5 | 105±4 | 127±7 | | | \$ |
| 3 | 91±12 | 97±8 | 114±15 | 150±9 | | | \$ |
| 4 | 95±8 | 126±14 | 139±18 | 111±16 |  |  |  |
| 5 | 85±6 | 96±3 | 94±13 | 64±4 | | | \$ |
| 6 | 107±5 | 107±7 | 132±14 | 114±5 |  |  |  |

Interaction effects of chemotherapy and sex: \$ p < 0.05: in weeks 2 and 3, chemotherapy-treated female mice ate more than control female mice. In week 5, chemotherapy-treated female mice ate less than control female mice and less than chemotherapy-treated male mice.

There were no significant treatment effects on heart and liver weights (Fig 1C). Chemotherapy increased spleen weight only in females (p<0.05), while, regardless of treatment, there was a sex effect on kidney weights, as females had significantly smaller kidneys compared to males (p<0.01). There was a main effect of chemotherapy on fat and muscle weights because irrespective of sex, chemotherapy treatment reduced the weights of the adipose tissue (Fig 1C, p<0.0001), gastrocnemius (Males: –26%, Females: –17%) and tibialis anterior (Males: –29%, Females: –24) skeletal muscles.

These effects on gastrocnemius muscle and adipose tissue were observed even in tissue raw weights (Table 3). There was also a main effect of sex on gastrocnemius muscle as regardless of treatment groups, females had greater weight compared to males (Fig 1D, p<0.05). For the quadriceps, chemotherapy-treated males experienced significant muscle loss compared to control males (Fig 1D, Table 3). There was a main effect of sex on raw liver, heart and kidney weights: irrespective of treatment groups, males had greater organ weights (Table 3). Because treatment and sex effects were observed in the gastrocnemius muscle, all muscle-based analyses were done in this muscle. There was a main effect of chemotherapy on the abundance of the myofibrillar protein myosin heavy chain-1 (MyHC-1) in that irrespective of sex, chemotherapy-treated groups had lower levels (p<0.01). Interestingly, compared to control males, chemotherapy-treated males exhibited reduced troponin and tropomyosin abundance (S1 Fig, p<0.001).

**Table 3.** Mean raw muscle, tissue and organ weights (mg)

|  | Control |  | Drug |  | Statistics |  |  |
| --- | --- | --- | --- | --- | --- | --- | --- |
| Tissue | Male | Female | Male | Female | Treatment | Sex | Interaction |
| Gastroc | 192±9 | 161±3 | 137±6 | 127±3 | *** |  |  |
| TA | 55±4 | 401±3 | 36±3 | 33±3 | | | \$\$ |
| Quad | 177±13 | 150±10 | 137±8 | 118±8 | | | \$ |
| Liver | 1460±107 | 1144±82 | 1464±50 | 1047±85 |  | # |  |
| Heart | 280±15 | 163±4 | 267±19 | 181±15 |  | ## |  |
| Spleen | 118±12 | 111±7 | 283±93 | 149±8 |  |  |  |
| Kidney | 414±15 | 212±12 | 486±67 | 191±10 |  | ## |  |
| Fat | 1392±148 | 1201±107 | 488±57 | 391±82 | **** |  |  |

Main effect of chemotherapy: \*\*\* p < 0.001, \*\*\*\* p < 0.0001; main effect of sex: # p < 0.05, ## p < 0.01; interaction effects of chemotherapy and sex: \$ p < 0.05, \$\$ p < 0.01. \$\$ Tibialis anterior (TA): male control have greater TA weight than female control and chemotherapy-treated male; \$ Quadriceps (Quad): chemotherapy reduced muscle weight only in male. Gastroc, gastrocnemius muscle.

### Chemotherapy-treated Males Exhibit Worsened Insulin Tolerance and Abundance of Ubiquitinated Proteins

During the ITT test, at weeks 3 and 6, there was no effect of chemotherapy on glucose AUC but there was a main effect of sex as glucose AUC was lower in females compared to males irrespective of treatment (Fig 2A-D). Also, at week 6, chemotherapy-treated males had higher glucose AUC than the male controls (Fig 2C, D, p<0.05). In mice that were fasted 3 h prior to sacrifice, there was a main effect of chemotherapy in that in gastrocnemius muscle, chemotherapy reduced the phosphorylation of AKT^ser473^, S6K1^thr389^ and S6^ser235/236^, consistent with decreased protein synthesis (measured by SuNSET analyses) in chemotherapy-treated mice (Fig 2E–G, p<0.05). Chemotherapy treatment did not affect total levels of AKT, S6K or S6 (S2 Fig) in both sexes. There were no main effects of chemotherapy or sex on the phosphorylation of FoxO3a^ser253^, abundance of MuRF1 (a muscle ubiquitin protein ligase) nor on ubiquitinated proteins, although chemotherapy-treated males exhibited increased abundance of ubiquitinated proteins compared to control males (Fig 2H – J).

**Fig 2:**
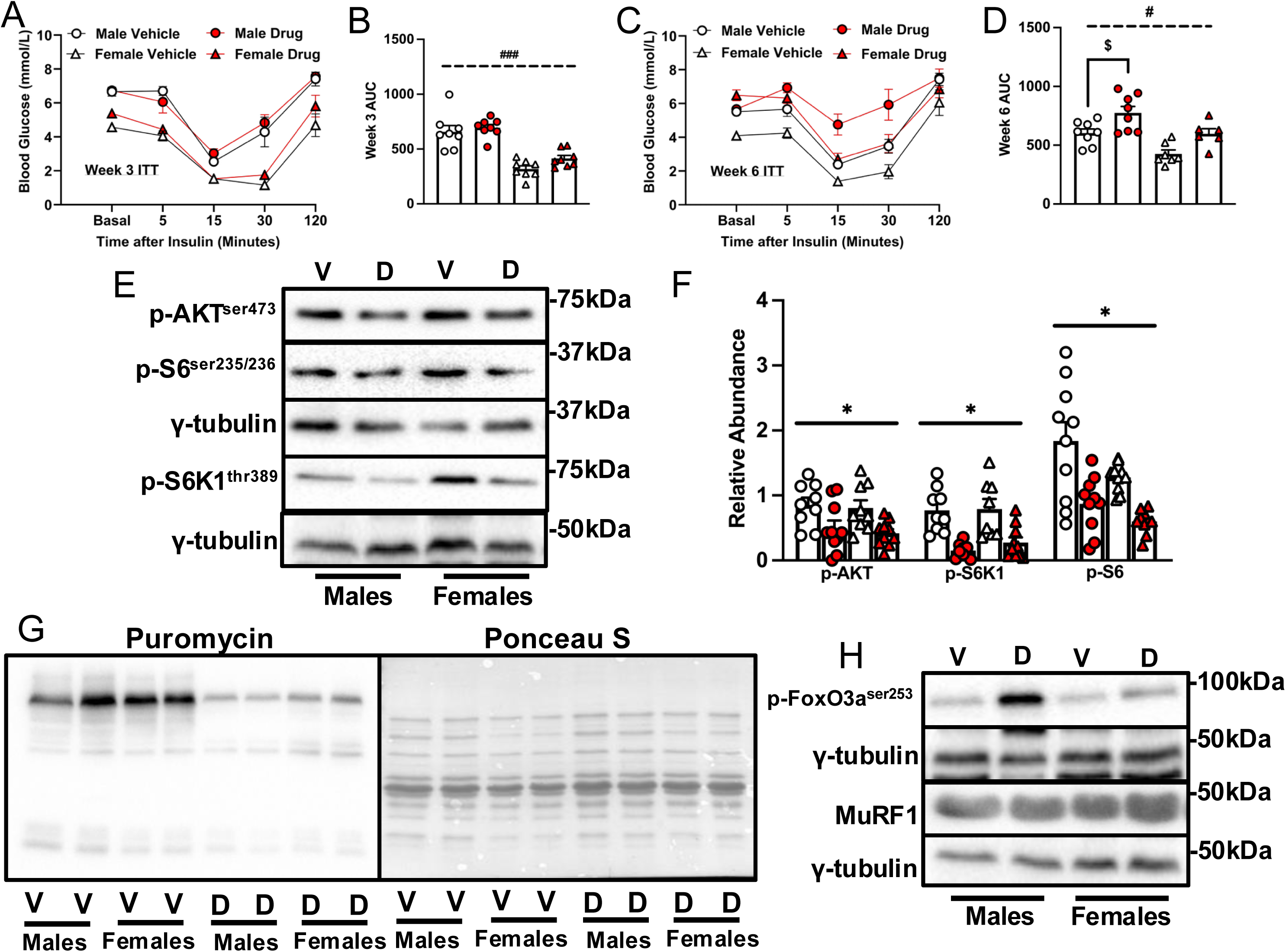
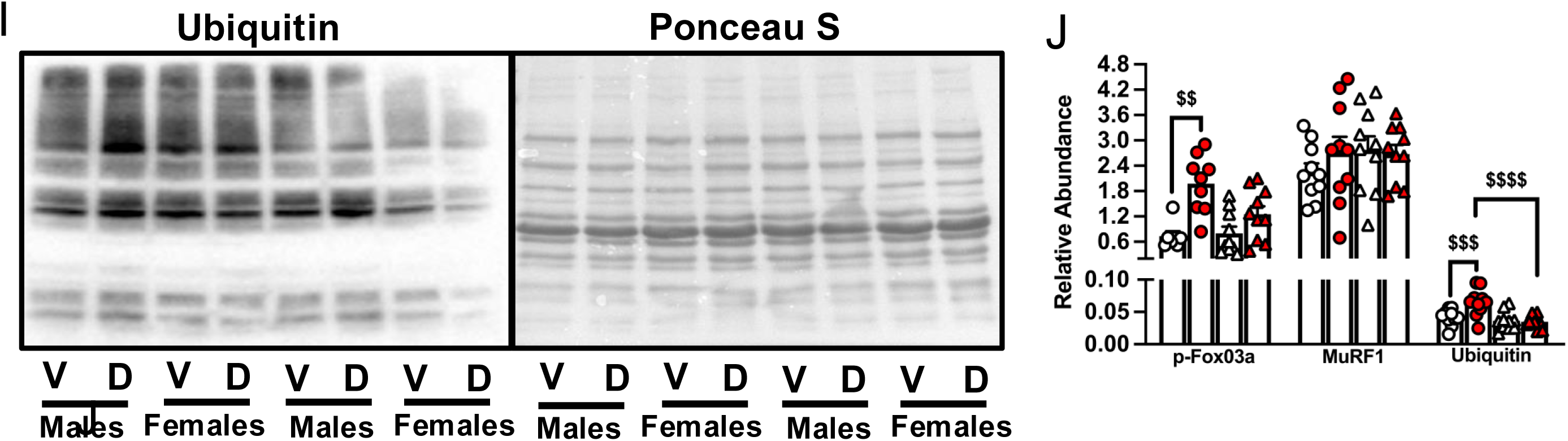
Worsened Insulin Tolerance and Abundance of Ubiquitinated Proteins in Old Male Mice in Response to Chemotherapy Treatment. Mice were treated as indicated in the legend to Fig 1. At least 24 h from the last chemotherapy dose, mice were starved for 6 h and underwent ITT by subcutaneous insulin (0.75units/kg) administration. Blood glucose curves and AUC are shown (A – D). Immunoblotting (E) and quantified data for p-AKTser473, p-S6K1thr389 and p-S6ser235/236 (F) are shown. Twenty-four h after their last chemotherapy dose and 3 h prior to euthanasia, protein synthesis was measured via the SUnSET analysis. Thirty minutes prior to euthanasia, mice were injected with 0.040μmol/g bodyweight of puromycin. Following euthanasia, tissues were collected and muscle proteins were blotted with an anti-puromycin antibody and corrected to their respective ponceau S staining (G). Immunoblotting (H, I) and quantified data for p-FoxO3Aser253, MuRF1 and ubiquitinated proteins (J) are shown. All anabolic and catabolic signals were measured in the gastrocnemius. Data are mean ± SE, n = 8 – 10. Main effect of chemotherapy: * p < 0.05, main effect of sex: # p < 0.05, ### p < 0.001, interaction effect of chemotherapy and sex: $ p < 0.05, $$ p < 0.01, $$$ p < 0.001, $$$$ p < 0.0001. Data were analyzed using a two-way ANOVA followed by a Tukey’s post hoc test. V, vehicle; D, drug (chemotherapy); AUC, area under curve; p, phosphorylation; h, hours; ITT, insulin tolerance test.

### Chemotherapy-treated Males Exhibit Increased Total Plasma but Reduced Muscle BCAA

Chemotherapy increased plasma levels of leucine in both sexes (main effect, p<0.001). Chemotherapy-treated males had higher total plasma BCAA levels compared to control males. This was not observed in females (Fig 3A, p<0.01). Also, only males showed decreased total plasma branched-chain keto acid (BCKA) levels following chemotherapy relative to the control group (Fig 3B, p<0.05). There was no main effect of chemotherapy or sex on gastrocnemius intracellular BCAA levels. However, there was a main effect of sex on valine in that the values were lower in females compared to males. Chemotherapy reduced muscle intracellular concentrations of leucine, valine, and total BCAA only in males (p<0.0001). For isoleucine, chemotherapy reduced its level only in females (Fig 3C, p<0.05). There was no main effect of chemotherapy or sex on muscle BCKA, but females treated with chemotherapy had reduced muscle KIV and total BCKA levels compared to female controls (Fig 3D, p<0.05). There were no significant treatment effects on glutamate (Fig 3E,) or on the other amino acids measured (S3 Fig). There was a main effect of chemotherapy on LAT1 level in that regardless of sex, there was reduced muscle abundance of LAT1 in chemotherapy-treated groups. Additionally, control females had higher LAT1 abundance compared to control males (Fig 3F). There was no treatment or sex effect on SNAT1 expression.

**Fig 3:**
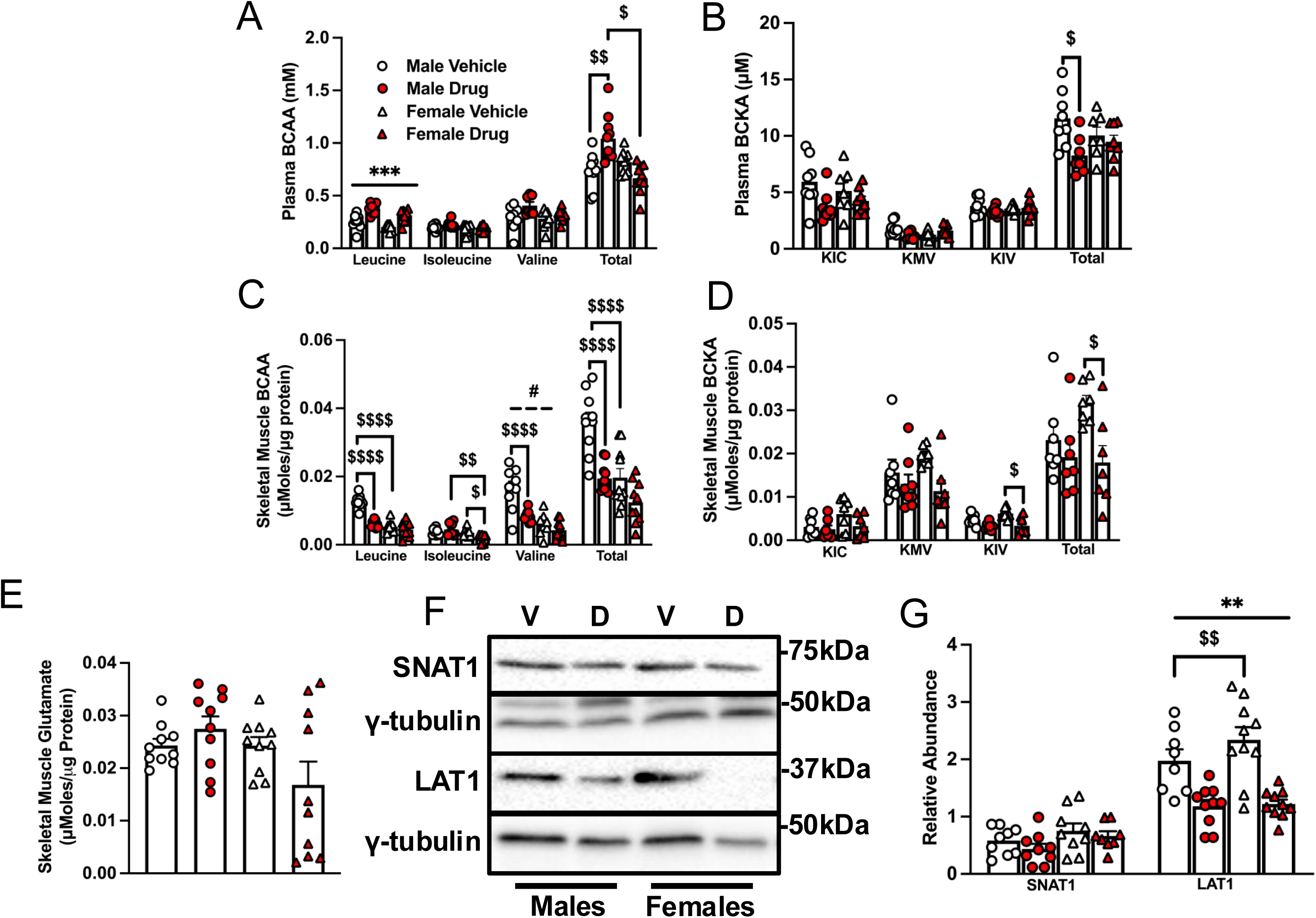
Chemotherapy-treated Old Males Exhibit Increased Total Plasma but Reduced Muscle BCAA. Mice were treated as indicated in the legend to Fig 1. Plasma BCAA (A) and BCKA (B), as well as gastrocnemius levels of the BCAA (C), BCKA (D) and glutamate (E) were measured by HPLC. Immunoblotting (F) and quantified data for SNAT1 and LAT1 (G) are shown in the gastrocnemius. Data are mean ± SE, n = 7 – 10. Main effect of chemotherapy: ** p < 0.01, *** p < 0.001; main effect of sex: # p < 0.05; interaction effects of chemotherapy and sex: $ p < 0.05, $$ p < 0.01, $$$$ p < 0.0001. Data were analyzed using a two-way ANOVA followed by a Tukey’s post hoc test. V, vehicle; D, drug (chemotherapy); BCAA, branched-chain amino acid; BCKA, branched-chain α ketoacids; HPLC, High pressure liquid chromatography; KIC, 2-keto-isocaproate/4-methyl-2-oxopentanoic acid; KMV, α-keto-β-methylvaleric acid/3-methyl-2-oxopentanoate; KIV, 2-keto-isovalerate/3-methyl-2-oxobutanoic acid.

### Chemotherapy Increases Measures of Skeletal Muscle BCAA Catabolism in Males

A simplified schematic of BCAA catabolism in skeletal muscle is shown in S4 Fig. In the gastrocnemius, there were no significant main effects of chemotherapy or sex on the abundance of BCAT2, BCKD and branched-chain α-ketoacid dehydrogenase kinase (BDK) (Fig 4A, B). However, chemotherapy reduced p-BCKD-E1α^ser293^ only in males (p<0.01), consistent with chemotherapy-induced increase in BCKD activity being observed in males but not in females (Fig 4A–C, p<0.05). Interestingly, there was a main effect of sex for p-BCKD-E1α^ser293^ in that irrespective of treatment groups, females had higher values compared to males (Fig 4A-B, p<0.0001).

**Fig 4:**
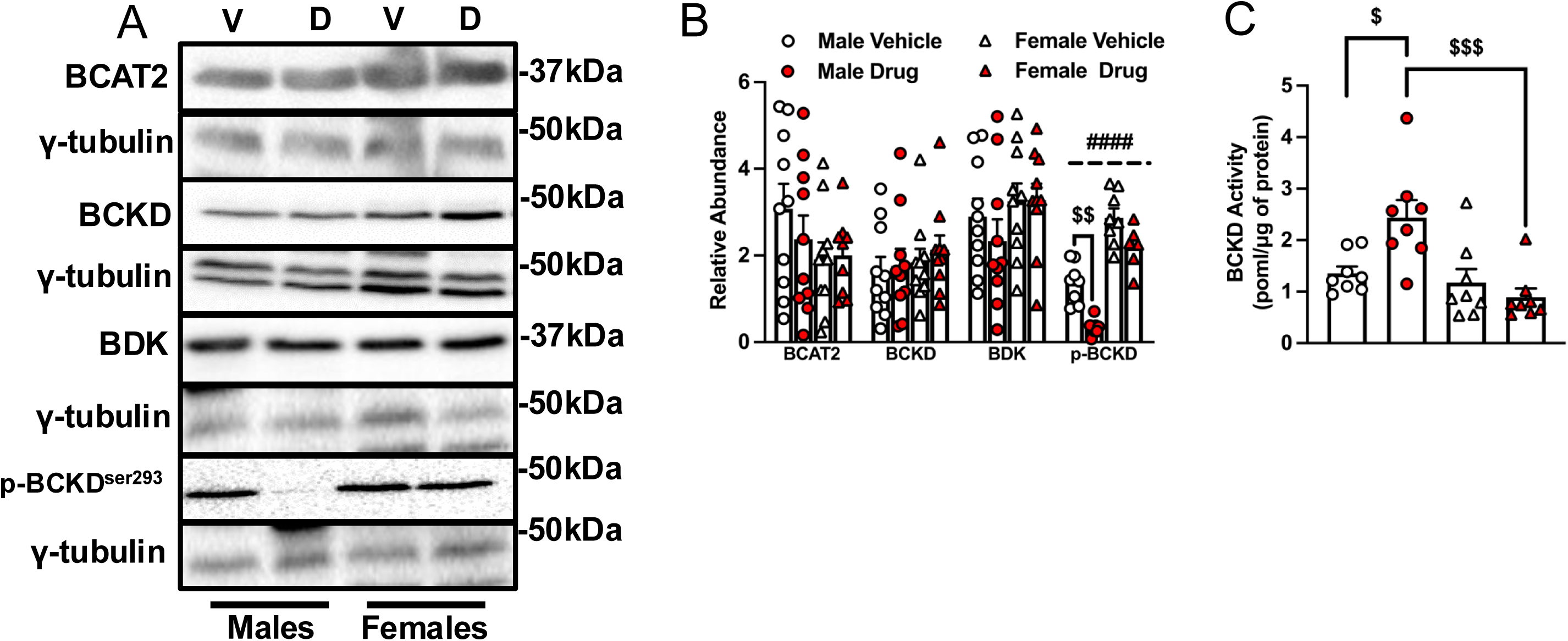
Measures of Skeletal Muscle BCAA Catabolism are Increased by Chemotherapy in Old Male Mice. Mice were treated as indicated in the legend to Fig 1. Key enzymes involved in BCAA catabolism were immunoblotted (A) and quantified data for BCAT2, BCKD-E1α, BDK and phosphorylated BCKD-E1αser293 (B) in the gastrocnemius are shown. Skeletal muscle BCKD activity was measured from the release of ^14^CO_2_ from ^14^C labelled valine (C). Data are mean ± SE, n = 7 – 10. Main effect of sex: #### p < 0.0001; interaction effects of chemotherapy and sex: $ p < 0.05, $$$ p < 0.001. Data were analyzed using a two-way ANOVA followed by a Tukey’s post hoc test. V, vehicle; D, chemotherapy drug; BCKD, branched-chain α-keto acid dehydrogenase complex; BCAA, branched-chain amino acid; BDK, branched-chain α-ketoacid dehydrogenase kinase.

### Chemotherapy Treatment Increases Indicators of BCAA Oxidation in the Liver of Old Mice of Either Sex

There were no main effects of chemotherapy or sex on liver intracellular BCAA or BCKA levels (Fig 5A-B). However, in females, chemotherapy reduced liver KMV, KIV and total BCKA levels (Fig 5B). There were no treatment effects on liver protein expression of LAT1, BCKD and BDK (Fig 5C, D), but there was a main effect of chemotherapy on p-BCKD-E1α^ser293^, with the values being lower in chemotherapy-treated groups irrespective of sex. This finding is consistent with a main effect of chemotherapy on liver BCKD activity: irrespective of sex, chemotherapy-treated groups had higher activity (Fig 5C – E).

**Fig 5:**
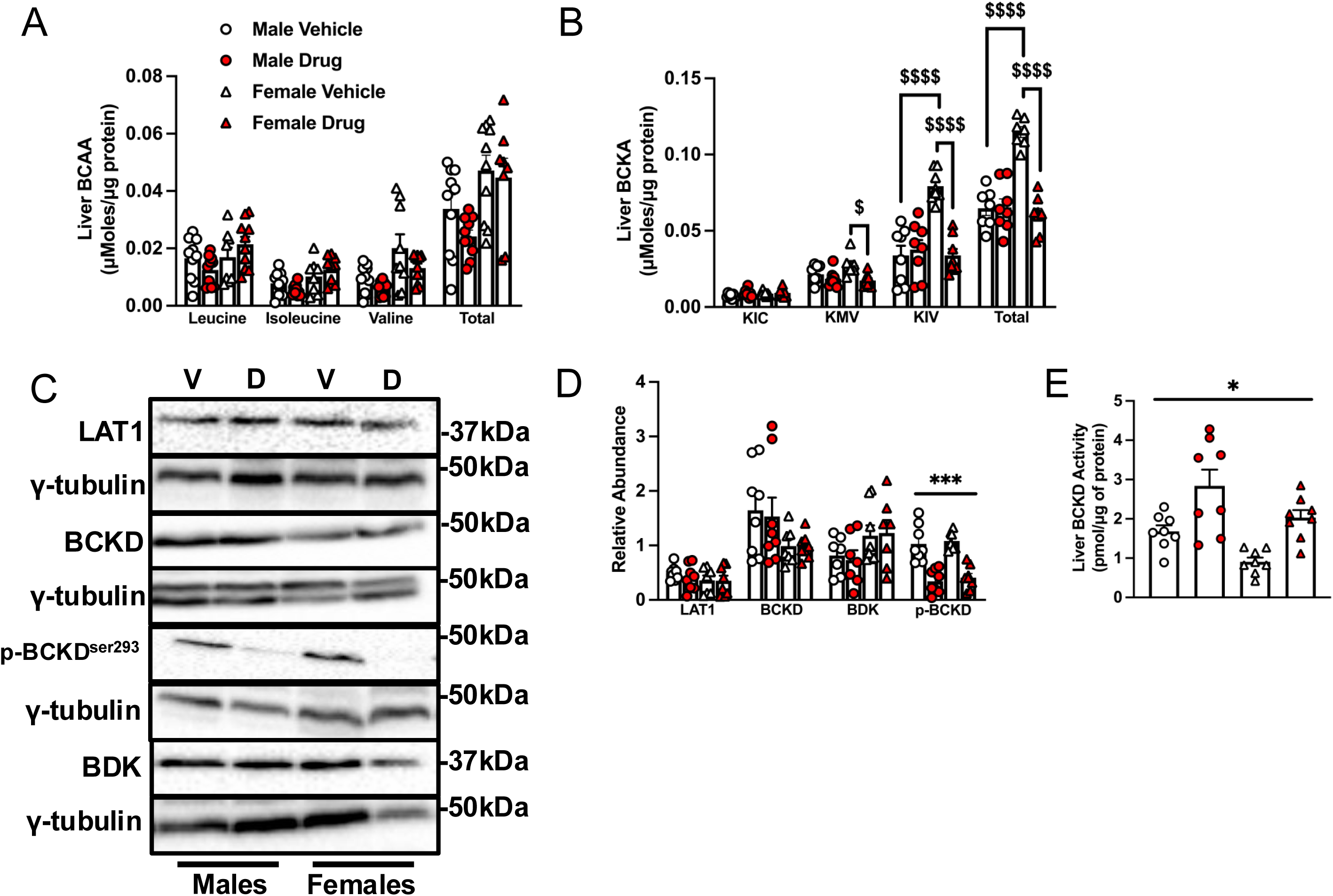
Chemotherapy Treatment Increases Indicators of BCAA Oxidation in the Liver of Both Sexes. Mice were treated as indicated in the legend to Fig 1. Liver samples were homogenized and the concentrations of the BCAA (A) and BCKA (B) were measured by HPLC. Immunoblotting (C) and quantified data for liver LAT1, BCKD-E1α, BDK and p-BCKD-E1αser293 (D) are shown. Liver BCKD activity was measured (E). Data are mean ± SE, n = 7 – 10. Main effect of chemotherapy: * p < 0.05, *** p < 0.001; interaction effects of chemotherapy and sex: $ p < 0.05, $$ p < 0.01, $$$$ p < 0.0001. Data were analyzed using a two-way ANOVA followed by a Tukey’s post hoc test. V, vehicle; D, chemotherapy drug; BCAA, branched-chain amino acid; HPLC, high pressure liquid chromatography. KIC, 2-keto-isocaproate/4-methyl-2-oxopentanoic acid; KMV, α-keto-β-methylvaleric acid/3-methyl-2-oxopentanoate; KIV, 2-keto-isovalerate/3-methyl-2-oxobutanoic acid; BCKD, branched-chain α-keto acid dehydrogenase complex; BDK, branched-chain α-ketoacid dehydrogenase kinase.

### Gastrocnemius Muscle Weight is Positively Correlated with Muscle Intracellular BCAA Levels, but not BCKD Activity

Positive correlations were found between total BCAA and gastrocnemius weight (Fig 6A) and LAT1 expression (Fig 6B), but not BCKD activity (Fig 6C). Gastrocnemius muscle weight was positively correlated with BCKD activity (Figure 6D) and LAT1 (Fig 6E).

**Fig 6:**
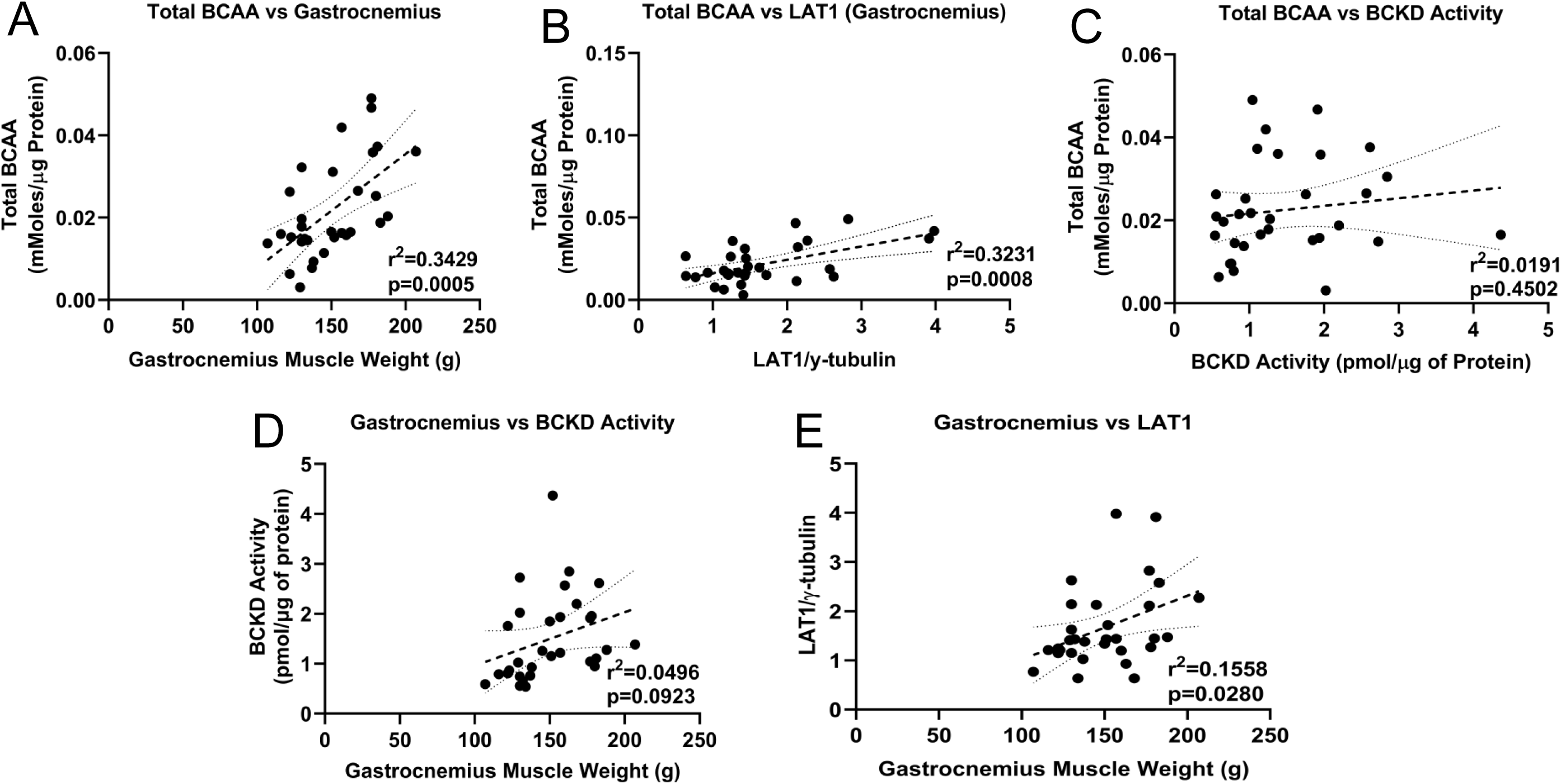
**Gastrocnemius Muscle Weight is Positively Correlated with Muscle Intracellular BCAA Levels**. Correlations between gastrocnemius muscle total intracellular BCAA concentrations and muscle weights (A), LAT1 expression (B), and skeletal muscle BCKD activity (C). Correlations between muscle weight and BCKD activity (D), and LAT1 (E) are also shown. Data were analyzed using linear regression and 95% confidence intervals are denoted. These charts are drawn from the data in Fig 1, 3 and 4. BCAA: Branched-chain amino acid; BCKD: branched-chain α-keto acid dehydrogenase complex.

## Discussion

Here, we report novel sex-related alterations in whole-body BCAA metabolism/handling following chemotherapy-induced cachexia in aged mice. In response to 6-week chemotherapy drug administration, we observed worsened insulin tolerance and greater loss of skeletal muscle weight, along with higher abundance of muscle ubiquitinated proteins in males. There was a main effect of chemotherapy on the abundance of LAT1, the canonical BCAA transporter, in that LAT1 level was reduced following chemotherapy in both sexes. However, there were chemotherapy-sex interactions in some of the other key measures of BCAA catabolism in chemotherapy-treated animals. For example, only chemotherapy-treated males showed decreases in skeletal muscle BCAA concentrations and p-BCKD-E1α^ser293^, corresponding to higher muscle BCKD activity. Also, in response to chemotherapy, muscle and liver BCKA levels were reduced only in females. Our findings of positive correlations between muscle mass and measures of BCAA metabolism suggest that sex-related alterations in BCAA catabolism may contribute to cachexia in older individuals.

Findings of sex differences in cachexia severity and measures of BCAA catabolism could be related to various factors, including differences in insulin sensitivity (58,59), hormonal profile (reviewed in (60)), variations in fiber type composition of muscles in males vs females (60,61), drug pharmacokinetics (62), and protein metabolism (63). Regarding insulin sensitivity and in line with findings from our study, male individuals show greater prevalence and development of insulin resistance (64). Sex differences in hormone levels, such as those seen for insulin (65,66), estrogen (67,68) and adiponectin (69) could account for these changes, however, we did not measure these here. While circulating levels of estrogen are reduced with aging (70), measurable levels can be seen in rodents as old as 20 months ((71,72)). Also, sex-dependent age-related differences in free estrogen and estrogen signaling (73,74) can explain some of the differences that we observed. In addition, in our study, high circulating levels of the BCAA, which is associated with insulin resistance (75), was only found in chemotherapy-treated males. For drug pharmacokinetics, female individuals show higher blood concentrations (76) and lower clearance rates (77) for the majority of FDA approved chemotherapy drugs, leading to higher hospitalizations due to adverse effects in females (76). Others have reported that compared to males, females have slower clearance of 5FU, one of the components of FOLFIRI (78,79).

We have shown that in young mice treated with chemotherapy, females experience greater body and muscle weight loss, consistent with greater loss of mTORC1 signalling (80). Here, in aged mice, we found a more severe loss of muscle weight in males following chemotherapy, a finding that is different from Huot et al’s (81). The difference in our findings might be related to the fact that Huot et al studied a different chemotherapy regimen and used a different mouse strain (C57BL/6J). Greater male muscle weight loss in our study was not due to sex differences in mTORC1 signalling, but occurred alongside greater insulin resistance and ubiquitinated proteins. These findings suggest that higher catabolism likely contributed to the greater skeletal muscle loss in males. Higher ubiquitinated proteins in males could be related to the fact that testosterone, a hormone associated with muscle mass loss in older males (82–84), can repress the expression of muscle ubiquitin ligases (85). However, protein expression of MuRF1, the E3 ligase responsible for the majority of protein ubiquitination in skeletal muscle (86), was not affected by chemotherapy treatment in either sex. Therefore, it is possible that the catabolic effects of aging on MuRF1 expression were already significant (87,88) and thus could not be worsened by chemotherapy. Nonetheless, in line with the fact that ubiquitin proteasome activity is higher in males (89,90), chemotherapy-induced changes in other E3 ligases that are increased in aging (88,91) but not measured here, such as atrogin-1 and parkin, could account for the higher ubiquitinated proteins in males.

Interventions that attenuate muscle wasting are at the forefront of cachexia studies. Due to their anabolic properties (92), we were interested in the effect of chemotherapy on the BCAA, as supplementation with these amino acids are ineffective in managing cachexia (34–37). In healthy individuals, BCAA increase protein synthesis and decrease protein breakdown (92). The fact that males have greater concentrations of skeletal muscle BCAA (41), implying that they could ‘afford’ to lose more, may in part explain our finding that chemotherapy reduced skeletal muscle BCAA only in males. Greater resistance to insulin in males could also explain this finding, due to the function of this hormone in regulating amino acid uptake (93). Although this finding could also be attributable to reductions in LAT1, there were no significant sex differences in the expression of this transporter, nor on SNAT1. Therefore, sex differences in BCAA catabolism (rather than in BCAA transport) may account for the changes in amino acid levels. For that reason, it would be interesting to study whether sex defences exist in the efficacy of BCAA supplementation in treating cachectic cancer patients.

While a previous study identified downregulation of proteins involved in amino acid metabolism in FOLFIRI-treated mice (39), that study investigated only male animals and did not identify how specific enzymes involved in the catabolism of the BCAA were affected. In our study, it was only in male mice that chemotherapy reduced p-BCKD-E1α^ser293^ and increased activity of this enzyme, implying greater skeletal muscle BCAA catabolism. These findings are consistent with data that showed that male individuals oxidize more amino acids for energy during substrate deficit conditions (94). Interestingly, there was a main effect of sex on muscle phosphorylation of BCKD-E1α^ser293^ in that irrespective of treatment, females had higher values. This would imply reduced BCKD activity and therefore reduced BCAA oxidation. This suggests that the BCKA, rather than being oxidized, are instead transaminated by BCAT2 back to the BCAA (28). This reasoning is consistent with findings of chemotherapy-induced reductions in muscle BCKA and, although not statistically significant, the tendency for glutamate to be lower in chemotherapy treated females relative to the corresponding drug control. Minimal changes in plasma BCKA and reduced liver BCKA in response to drug treatment in female further support this reasoning, as the keto acids can travel back to the skeletal muscle to restore the BCAA pool. These sex-dependent patterns of changes following chemotherapy suggests maintenance of the BCAA pool to protect against atrophy in female but not in male mice. The effect of chemotherapy on transcription factors that are known to regulate BCAA metabolism, such as peroxisome proliferator-activated receptor gamma coactivator 1-alpha (95) and Krueppel-like factor 15 (96) may also be responsible for the changes we observed.

A main limitation of this study is the fact that we did not include any intervention group. However, because studying the effects of chemotherapy drug cocktails in old mice is rarely done, it was not clear what kind of interventions would be appropriate and therefore, findings from our study would be invaluable in designing future interventional studies. Another limitation is that we did not measure the effect of chemotherapy on adipose tissue, as this tissue too is a main site of BCAA metabolism (97,98). Also, BCAA metabolism is often studied in tumour cells (99,100), but rarely in host tissues, therefore, we studied clinically relevant anti-cancer drugs (101) at doses that do not exceed clinical levels (38), as a model to induce cachexia and probe sex differences in skeletal muscle BCAA metabolism in healthy mice. Related, while we have reported reduced muscle intracellular BCAA in response to chemotherapy, it is not known if chemotherapy and/or tumour implantation specifically alter intestinal absorption of BCAA. This is an important question that is outside of the scope of this work. Cancer cells, including breast and liver cancers, can alter BCAA metabolism to favour their growth (99,100). BCAA-focussed nutritional interventions in cachectic cancer patients must take this into consideration. Because tissue samples were collected ∼24h after the last drug administration, an acute effect of the drug cocktail could not be ruled out. However, the fact the main effect of chemotherapy and chemotherapy-sex interactions on body weight became more pronounced toward the end of the study (Fig 1A and Table 1) points to a cumulative chronic effect of the chemotherapy drugs. Nevertheless, studies in which animals are euthanized at different times after initiation of chemotherapy can help to identify acute vs chronic effects of the drug cocktail. Also, developmental studies in which animals are studied at different ages can help to characterize the ontogeny of age effects on BCAA catabolism in response to chemotherapy. While we have used chemotherapy drugs as agents that can induce cachexia and alter BCAA metabolism, clinically, chemotherapy drugs are used to treat cancer. Therefore, future studies in which tumour-bearing animals are treated with chemotherapy drugs will be useful to identify the combined effects of these two factors in settings that are clinically relevant.

In summary, compared to female animals, aged male mice exhibited greater loss of skeletal muscle weight and intracellular BCAA concentrations following chemotherapy. Given the strong correlations between muscle BCAA and muscle mass (Fig 6), altered muscle BCAA metabolism may be a reason for the relative ineffectiveness of nutritional interventions to mitigate cachexia, especially in older individuals. Observations from this study also suggest that during cachexia, BCAA nutritional support may be less effective in older males, as increasing BCAA supply, which theoretically should support increased anabolism, would instead lead to increased catabolism and oxidation of these amino acids. These novel findings also provide a mechanistic viewpoint to support previous reports highlighting greater prevalence of cachexia in males (102,103) and point to a need for a consideration of sex-specific approaches to manage cachexia.

## Data availability

Data will be made available upon reasonable request.

## Acknowledgements

The authors thank the Muscle Health Research Centre at York University for use of the HPLC and Dr. Paluzzi Lab for the use of their imaging systems.

## Author contributions

SM and OAJA designed the experiments. SM and GM performed the experiments. SM analyzed the samples and drafted the initial version of the manuscript. OAJA and GM reviewed and edited the manuscript. SM, GM and OAJA approve the final version of the manuscript.

## Ethics statement and experimental approval

All mice experiments were approved by the York University Animal Care Committee and were conducted within the guidelines of the Canadian Council on Animal Care.

## Supporting information

**S1 Fig. Related to Manuscript Fig 1. Myofibrillar proteins are decreased following chemotherapy treatment.** Male and female (18±2 months of age) CD2F1 mice were treated with either vehicle (males: white circle, females: white triangle; 3.8% DMSO in saline) or a chemotherapy drug cocktail (males: red circle, females: red triangle; 50mg/kg 5FU, 90mg/kg Leucovorin, 24mg/kg CPT11; Drug) twice per week for 6 weeks. Immunoblots **(A)** and quantified blots for the myofibrillar proteins **(B)**. Data are mean ± SE, n = 8 – 10. Main effect of chemotherapy: ** p < 0.01, interaction effects of chemotherapy and sex: $$ p < 0.05, $$$ p < 0.001, $$$$ p < 0.0001. For example, chemotherapy reduced troponin only in males. Data were analyzed using a two-way ANOVA followed by a Tukey’s post hoc test. V, vehicle; D, chemotherapy drug.

**S2 Fig, related to Manuscript Fig 2. Total protein levels of signaling proteins are not affected by chemotherapy treatment.** Male and female (18±2 months of age) CD2F1 mice were treated as described in supplementary figure 1. Immunoblots are shown for the total levels of AKT, S6 and S6K1 in the gastrocnemius **(A)**. V, vehicle; D, chemotherapy drug.

**S3 Fig. Related to Manuscript Fig 3. Minimal changes were found for other amino acids measured.** Male and female (18±2 months of age) CD2F1 mice were treated as described in supplementary figure 1. Amino acids were measured by HPLC in the gastrocnemius **(A)**. Data are mean ± SE, n = 8 – 10. Interaction effects of chemotherapy and sex: $ p < 0.05. Data were analyzed using a two-way ANOVA followed by a Tukey’s post hoc test.

**S4 Fig Related to Manuscript Fig 4. A simplified diagram of BCAA catabolism in skeletal muscle.** The BCAA are first reversibly transaminated by BCAT2. The resulting BCKA are then irreversibly decarboxylated by the BCKD complex, yielding isovaleryl-CoA, 2-methylbutyryl-CoA and isobutyryl-CoA. Each of these is then funneled into their respective metabolic pathways. BCAA, branched-chain amino acid ; BCKA, branched-chain α-ketoacids; BCAT, branched-chain aminotransferase; BCKD, branched-chain α-keto acid dehydrogenase complex; KIC, 2-keto-isocaproate/4-methyl-2-oxopentanoic acid; KMV, α-keto-β-methylvaleric acid/3-methyl-2-oxopentanoate; KIV, 2-keto-isovalerate/3-methyl-2-oxobutanoic acid. Re-drawn and modified from Mann et al (28).

**S1 Table. List of animal studies investigating cancer- and chemotherapy-induced cachexia**

**S2 Table. List of primary antibodies**

**S3Table. Daily food intake**

