## Supporting information for "Sex Differences in Cachexia Outcomes and Branched-Chain Amino Acid Metabolism Following Chemotherapy in Aged Mice"

### Supplementary figures for "Sex Differences in Cachectic Outcomes and Branched-Chain Amino Acid Metabolism following Chemotherapy in Old Mice"

Stephen Mora, Gagandeep Mann, Olasunkanmi A J Adegoke*

**S1 Table: Publications on Animal Studies Investigating Cancer- and Chemotherapy-Induced Cachexia**

| Sex / Animal | Age (Weeks) | Cancer | Chemotherapy | Body Weight Loss (~%) | Muscle Weight Loss (~%) | Ref |
|---|---|---|---|---|---|---|
| Male Wistar Rats | 3 | Walker 256 | -- | 10 | -- | (1) |
| Male Wistar Rats | 5 | AH-130 Hepatoma | -- | -- | GA: 13<br>EDL: 18<br>TA: 12 | (2) |
| Female Nude Mice | 5 | SEKI Human Melanoma | -- | 20 | -- | (3) |
| C57BL/6J Mice (no sex given) | 5-6 | Lewis Lung Carcinoma | -- | 25 | GA: 30 | (4) |
| Male CD2F1 Mice | 6 | C26 Adenocarcinoma | -- | Significant body weight change | Significantly smaller gain in muscle mass | (5) |
| Male Balb/c Mice | 6 | C26 Adenocarcinoma | -- | 10 | -- | (6) |
| Male Balb/c and C57BL/6J Mice | 6 | C26 Adenocarcinoma | -- | 20 | GA: 20<br>TA: 22 | (7) |
| Male and Female Balb/c Mice | 6 | C26 Adenocarcinoma | 5FU (50mg/kg) and oxaliplatin (6mg/kg), 3 injections | Cancer: 15<br>Cancer + Chemo: 22 | Cancer GA: 25<br>Cancer + Chemo GA: 40 | (8) |
| Male C57BL/6J Mice | 5-7 | Lewis Lung Carcinoma | -- | 25% less weight gained compared to control | -- | (9) |
| Male CD2F2 Mice | 6-7 | C26 Adenocarcinoma | -- | 10 | TA: 20 | (10) |
| CD2F1 Mice (no sex given) | 6-8 | C26 Adenocarcinoma | -- | 30 | TA: 40<br>GA: 40<br>QU: 50 | (11) |
| Male C57BL/6J Mice | 7 | Lewis Lung Carcinoma | -- | Significant Decrease | TA: ~30<br>EDL: ~40 | (12) |
| Male CD2F1 Mice | 7-8 | C26 Adenocarcinoma | Cisplatin (5mg/kg), 4 injections | Cancer: 5<br>Chemo: 15<br>Cancer + Chemo: 10 | Cancer GA: 15<br>Chemo GA: 15<br>Cancer + Chemo GA: 15 | (13) |
| Male CD2F1 Mice | 8 | C26 Adenocarcinoma | 5FU (50mg/kg), LEU (90mg/kg), CPT-11 (24mg/kg) for 5 weeks | Cancer: 30<br>Chemo: 15<br>Cancer + Chemo: 30 | Cancer: GA: 25<br>Cancer: TA: 20<br>Cancer: QU: 20<br>Chemo: GA: 15<br>Chemo: TA: 10<br>Chemo: QU: 15<br>Cancer + Chemo: GA: 40<br>Cancer + Chemo: TA: 25<br>Cancer + Chemo: QU: 40 | (14) |
| Male CD2F1 Mice | 8 | -- | 5FU (30mg/kg), LEU (90mg/kg), CPT-11 (24mg/kg) for 5 weeks | 10 | GA: 10<br>TA: 15<br>QU: 25 | (15) |
| Male CD2F1 Mice | 8 | C26 Adenocarcinoma | 5FU (30mg/kg), LEU (90mg/kg), CPT-11 (24mg/kg) for 5 weeks | Cancer: 13<br>Chemo: 15 | Cancer GA: 23<br>Cancer QU: 25<br>Chemo GA: 11<br>Chemo QU: 20 | (16) |

| | | | | | | |
|---|---|---|---|---|---|---|
| Male CD2F1 Mice | 8 | -- | 5FU (30mg/kg), LEU (90mg/kg), CPT-11 (24mg/kg) for 5 weeks | -- | GA: 10 | (17) |
| C57BL/6J Mice (no sex given) | 8 | -- | Cisplatin (2.5mg/kg), 9 injections | -- | GA: 10 | (17) |
| Female Sprague-Dawley Rats | 8 | -- | Doxorubicin (4mg/kg), 3 injections | 15 | SOL CSA: 25<br>EDL CSA: 35 | (18) |
| Male C57BL/6J Mice | 8 | Lewis Lung Carcinoma | -- | -- | TA: 30<br>EDL: 30 | (19) |
| Male C57BL/6J Mice | 8-9 | -- | Cisplatin (3m/kg), 4 injections | 15 | QU: 18<br>HL: 20 | (20) |
| Male C57BL/6J Mice | 6-10 | -- | Doxorubicin (15mg/kg), 1 injection | -- | All Muscles: 10 | (21) |
| Male Balb/c Mice | 8-10 | -- | Doxorubicin (2.5mg/kg) for 4 weeks | 10 | TA: 10<br>EDL: 13<br>SOL: 8 | (22) |
| Male Wistar Rats | 8-12 | Walker 256 | -- | 10 | -- | (23) |
| Male Wistar Rats | 8-12 | Walker 256 | -- | 2 | -- | (24) |
| Male C57BL/6J Mice | 9-10 | -- | Doxorubicin (4mg/kg), 4 injections | 10 | GA: 15<br>TA: 8<br>SOL: 5 | (25) |
| Male Sprague-Dawley Rats | 10 | -- | Doxorubicin (15mg/kg), 1 injection | 2 | EDL: 2 | (26) |
| Male C57BL/6J Mice | 10 | Lewis Lung Carcinoma | -- | 10 | GA: 10<br>SOL: 15 | (27) |
| Male Wistar Rats | 10 | Walker 256 | -- | Significant Decrease | GA: 25 | (28) |
| Male Rats: F344/NTacfBR | 10-12 | Methylcholanthrene Sarcoma | -- | 10 | -- | (29) |
| Male and Female C57BL/6J | 12 | Methylcholanthrene Sarcoma | -- | 20 | -- | (30) |
| Male C57BL/6J Mice | 12 | Lewis Lung Carcinoma | -- | 20 | QU: ~25<br>GA: ~25<br>TA: ~50 | (31) |
| Female Wistar Rats | 12 | AH-130 Hepatoma | -- | 5 | HL: 40 | (32) |
| Male Wistar Rats | 14 | -- | Doxorubicin (15mg/kg), 1 injection | Significant change (-30.05) from Control | EDL: 10 | (33) |
| Male and Female C57BL/6J Mice | 16 | -- | 5FU (30mg/kg), LEU (90mg/kg), CPT-11 (24mg/kg) for 9 weeks | 12 | Lean Mass: 15 | (34) |
| Male C57BL/6J Mice | 18-20 | Lewis Lung Carcinoma | -- | 5 | GA: 15<br>TA: 15 | (35) |
| Male C57BL/6J Mice | 20-28 | Lewis Lung Carcinoma | -- | 20 | -- | (36) |
| Male and Female C57BL/6J Mice | 8 (Yo) and 72 (Old) | -- | Cisplatin (2.5mg/kg), 9 injections | Yo Male: 13<br>Yo Female: 12<br>Old Male: 34<br>Old Female: 27 | Yo Male GA: 16<br>Yo Female GA:11<br>Old Male GA: 22<br>Old Female GA: 27 | (37) |
| Male Balb/c Mice | 8-16 (Yo) and 60-80 (Old) | C26 Adenocarcinoma | -- | Yo Male: 15<br>Old Male: 13 | Yo Male GA: 10<br>Old Male GA: 10 | (38) |

| | | | | | | |
|---|---|---|---|---|---|---|
| Male Lister Hooded Rats | -- | -- | Cisplatin (4mL/kg) | 6 | EDL: 8 | (39) |
| Male Wistar Rats | -- | AH-130 Hepatoma | -- | -- | GA: 10<br>SOL: 15 | (40) |
| Male C57BL/6J Mice | -- | Lewis Lung Carcinoma | -- | 5 | -- | (41) |
| Male Wistar Rats and Male C57BL/6J Mice | -- | AH-130 Hepatoma and C26 Adenocarcinoma | -- | Rat: 20<br>Mice: 15 | Rat GA: 20<br>Mice GA: 20 | (42) |
| Male Wistar Rats | -- | C26 Adenocarcinoma | -- | 20 | GA: 20<br>TA: 20 | (43) |
| Male Wistar Rats | -- | AH-130 Hepatoma | -- | 25 | GA: 25 | (44) |
| Male Wistar Rats | -- | AH-130 Hepatoma | -- | 30 | GA: 25 | (45) |

GA, Gastrocnemius; TA, Tibialis Anterior; QU, Quadriceps; SOL, Soleus; EDL, Extensor Digitorum Longus; HL, Hindlimb; 5FU, 5-fluorouracil; CPT-11, Irinotecan Hydrochloride; LEU, Leucovorin; CSA, Cross Sectional Area; Yo, young; --, not measured/information not provided; Chemo, Chemotherapy.

**S2 Table – Primary Antibodies**

| ANTIBODY | SOURCE | DILUTION | RRID | COMPANY |
|---|---|---|---|---|
| MyHC-1 | Mouse | 1:500 | AB_2147781 | Developmental Hybridoma (MF-20) |
| Troponin | Mouse | 1:400 | AB_2618103 | Developmental Hybridoma (JLT12) |
| Tropomyosin | Mouse | 1:400 | AB_2205770 | Developmental Hybridoma (CH-1) |
| p-FoxO3a$^{ser253}$ | Rabbit | 1:1000 | AB_2106674 | Cell Signalling Tech (#9466) |
| p-AKT$^{ser473}$ | Rabbit | 1:1000 | AB_2315049 | Cell Signalling Tech (#4060) |
| p-S6$^{ser235/236}$ | Rabbit | 1:1000 | AB_916156 | Cell Signalling Tech (#4858) |
| p-S6K1$^{thr389}$ | Rabbit | 1:1000 | AB_2269803 | Cell Signalling Tech (#9234) |
| SNAT1 | Rabbit | 1:1000 | AB_2799092 | Cell Signalling Tech (#36057) |
| p-BCKD-E1α$^{ser293}$ | Rabbit | 1:1000 | AB_2799176 | Cell Signalling Tech (#40368) |
| BCKD-E1α | Rabbit | 1:1000 | AB_2800155 | Cell Signalling Tech (#90198) |
| BCAT2 | Rabbit | 1:1000 | AB_10792411 | Protein Tech (#16417-1-AP) |
| MuRF1 | Rabbit | 1:1000 | AB_11232209 | Protein Tech (#55456-1-AP) |
| BDK | Rabbit | 1:1000 | AB_2548929 | Invitrogen (#PA5-31455) |
| LAT1 | Rabbit | 1:500 | AB_2635938 | Invitrogen (#PA5-50485) |
| γ-tubulin | Mouse | 1:10000 | AB_477584 | Sigma Aldrich (#T6557) |
| Puromycin | Mouse | 1:20000 | AB_2566826 | Sigma Aldrich (#MABE343) |
| Ubiquitin | Mouse | 1:500 | AB_628423 | Santa Cruz (#SC-8017) |

**S3Table. Daily food intake**

| | **Cont-Male** | | **Cont-Fem** | | **Drug-Male** | | **Drug-female** | | **Statistics** | | |
|---|---|---|---|---|---|---|---|---|---|---|---|
| Day | **Mean** | **SD** | **Mean** | **SD** | **Mean** | **SD** | **Mean** | **SD** | **Trt** | **Sex** | **Int** |
| D2 | 3.7 | 1.3 | 3.5 | 1.1 | 3.0 | 1.3 | 2.8 | 0.9 | | | |
| D3 | 3.5 | 0.8 | 2.5 | 1.0 | 2.4 | 0.9 | 1.6 | 0.5 | | | |
| **D4** | **3.2**a | 0.7 | **2.1**b | 0.4 | 2.5 | 1.0 | 1.9 | 0.5 | | | 0.005 |
| D5 | 3.1 | 0.5 | 2.5 | 0.1 | 2.7 | 0.6 | 1.8 | 0.0 | | .04 | |
| D6 | 4.0 | 1.2 | 2.8 | 0.4 | 3.4 | 0.9 | 2.3 | 0.7 | | .04 | |
| D7 | 4.0 | 1.2 | 2.6 | 0.7 | 3.4 | 0.9 | 2.3 | 0.7 | | .04 | |
| **D8** | 3.5 | 1.7 | 2.7 | 0.6 | 3.0 | 1.2 | 2.5 | 0.7 | | | |
| D9 | 3.5 | 0.8 | 3.4 | 1.2 | 2.9 | 1.0 | 2.2 | 0.2 | | | |
| D10 | 3.3 | 0.8 | 2.8 | 0.8 | 2.8 | 0.8 | 2.4 | 0.6 | | | |
| **D11** | $3.3^{a}$ | 0.9 | $3.1^{a}$ | 0.3 | $2.7^{a}$ | 0.7 | $1.8^{b}$ | 0.5 | | | .03 |
| D12 | 4.0 | 1.2 | $3.2^{a}$ | 0.7 | 3.7 | 0.9 | $4.5^{b}$ | 0.4 | | | .001 |
| D13 | 4.2 | 1.2 | 3.6 | 0.3 | 3.8 | 0.7 | 3.5 | 0.9 | | | |
| D14 | 4.1 | 1.2 | 3.6 | 0.3 | 3.9 | 0.7 | 3.5 | 0.9 | | | |
| **D15** | 3.8 | 0.7 | $3.3^{a}$ | 0.0 | 2.7 | 1.1 | $2.1^{b}$ | 0.5 | | | .008 |
| D16 | 4.4 | 2.0 | $4.0^{a}$ | 0.0 | 2.8 | 1.1 | $2.4^{b}$ | 0.0 | | | .0001 |
| D17 | 3.6 | 1.2 | 4.6 | 0.3 | 4.2 | 1.4 | 4.2 | 1.0 | | | |
| **D18** | 4.0 | 1.8 | 2.8 | 0.2 | 4.9 | 2.3 | 2.9 | 1.6 | | | |
| D19 | 3.5 | 1.3 | $3.5^{a}$ | 0.7 | 3.8 | 1.0 | $1.8^{b}$ | 0.4 | | | .0001 |
| D20 | 3.2 | 0.5 | $3.5^{a}$ | 0.6 | 3.6 | 0.7 | $1.9^{b}$ | 0.4 | | | .0001 |
| D21 | 3.2 | 0.5 | $3.3^{a}$ | 0.4 | 3.6 | 0.7 | $1.9^{b}$ | 0.4 | | | .0001 |
| **D22** | 3.5 | 0.8 | $4.3^{a}$ | 1.3 | 3.5 | 2.0 | $1.6^{b}$ | 0.2 | | | .0003 |
| D23 | 3.5 | 1.0 | $5.0^{a}$ | 1.7 | 3.5 | 1.5 | $2.4^{b}$ | 0.5 | | | .0004 |
| D24 | 4.5 | 1.4 | 4.6 | 0.8 | 4.1 | 1.5 | 4.0 | 0.8 | | | |
| **D25** | 3.6 | 0.6 | $3.5^{a}$ | 0.3 | 4.6 | 2.9 | $2.7^{b}$ | 0.7 | | | .03 |
| D26 | 3.8 | 1.1 | $3.5^{a}$ | 0.5 | 3.8 | 0.9 | $2.4^{b}$ | 0.2 | | | .006 |
| D27 | 4.0 | 1.1 | $3.1^{a}$ | 0.3 | 4.1 | 1.5 | $1.7^{b}$ | 0.9 | | | .003 |
| D28 | 4.1 | 1.1 | $3.1^{a}$ | 0.3 | 4.3 | 1.3 | $2.0^{b}$ | 0.5 | | | .001 |
| **D29** | 4.0 | 0.9 | $3.6^{a}$ | 1.0 | 4.3 | 2.0 | $1.4^{b}$ | 0.2 | | | .005 |
| D30 | 3.9 | 1.0 | 3.9 | 0.8 | 4.6 | 2.9 | 3.4 | 1.0 | | | |
| D31 | 4.0 | 0.7 | $4.4^{a}$ | 0.6 | 4.0 | 2.3 | $2.6^{b}$ | 0.4 | | | .0001 |
| **D32** | 3.3 | 1.1 | 2.5 | 0.1 | 2.6 | 1.2 | 2.7 | 0.4 | | | |
| D33 | 4.5 | 1.0 | $3.3^{a}$ | 0.0 | 4.1 | 1.2 | $2.7^{b}$ | 0.4 | | | .04 |
| D34 | 4.2 | 0.7 | $3.3^{a}$ | 0.0 | 4.6 | 1.0 | $2.8^{b}$ | 0.3 | | .04 | .003 |
| D35 | 3.9 | 0.8 | $3.3^{b}$ | 0.0 | 4.6 | 1.0 | $2.8^{b}$ | 0.2 | | | .004 |

S3 Table. Related to Manuscript Fig 1 and Table2. Male and female (18±2 months of age) CD2F1 mice were treated with either vehicle (control (Cont); 3.8% DMSO in saline) or a chemotherapy drug cocktail (Drug; 50mg/kg 5FU, 90mg/kg Leucovorin, 24mg/kg CPT11) twice per week for 6 weeks. There was no main effect of chemotherapy on any of the study days. On some days, males ate more than females (sex effect). The interaction effects seen on some days were largely from chemotherapy-treated female mice consuming less than female control. Day 1 was the day of first chemotherapy treatment. Other treatment days are in bold, along with day 36 (not shown). P values of statistical analyses are shown in the column “Statistics.” Cont Male: control (vehicle-treated) male; Cont-Fem, control female; Trt, treatment; Int, interaction effects of chemotherapy and sex; SD, standard deviation.

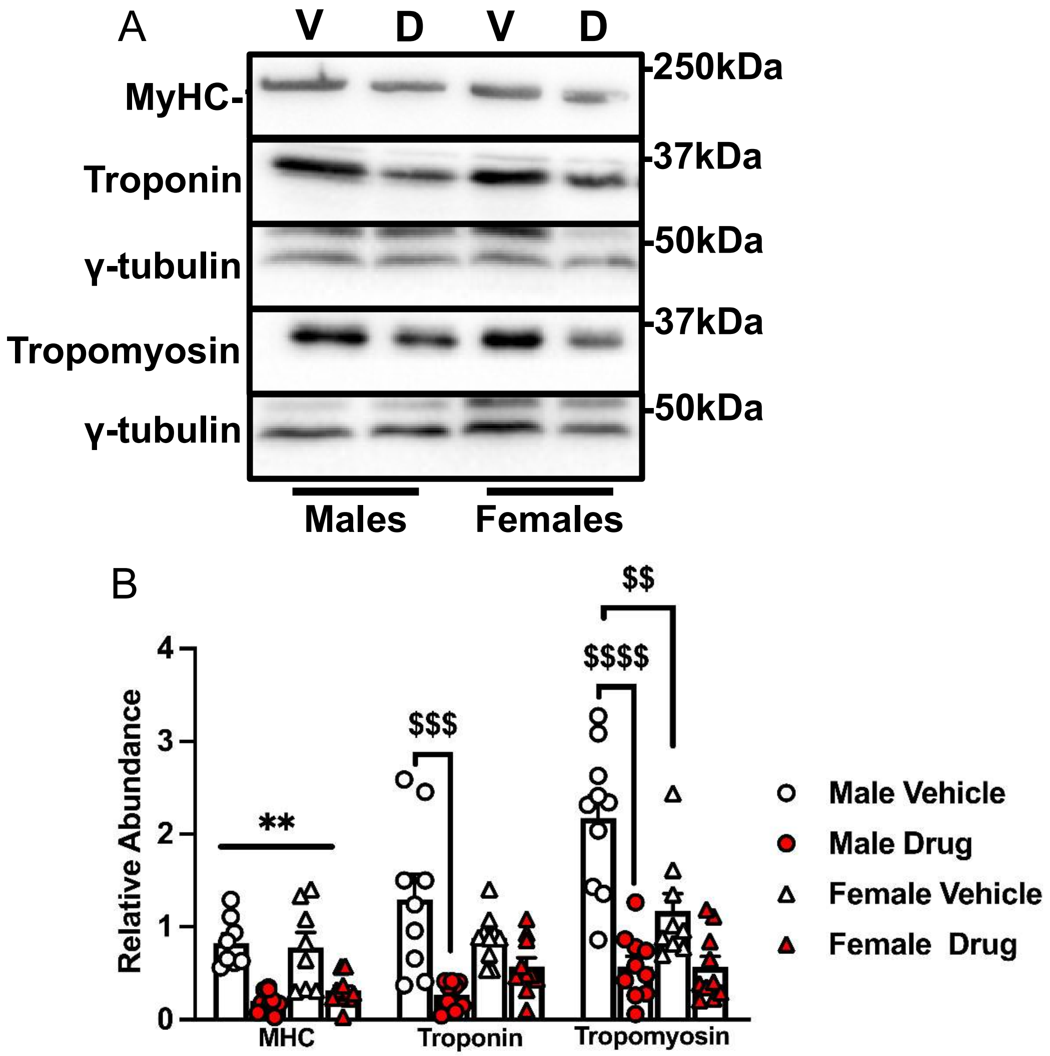


**S1 Fig. Related to Manuscript Fig 1. Myofibrillar proteins are decreased following chemotherapy treatment.** Male and female (18±2 months of age) CD2F1 mice were treated with either vehicle (male: white circle, female: white triangle; 3.8% DMSO in saline) or a chemotherapy drug cocktail (male: red circle, female: red triangle; 50mg/kg 5FU, 90mg/kg Leucovorin, 24mg/kg CPT11; Drug) twice per week for 6 weeks. Immunoblots **(A)** and quantified blots for the myofibrillar proteins **(B)**. Data are mean ± SE, n = 8 – 10. Main effect of drug: ** $p < 0.01$, effect of drug and sex interaction: $$ $p < 0.05$, $$$ $p < 0.001$, $$$$ $p < 0.0001$. For example, chemotherapy reduced troponin only in male. Data were analyzed using a two-way ANOVA followed by a Tukey's post hoc test. V, vehicle; D, drug.

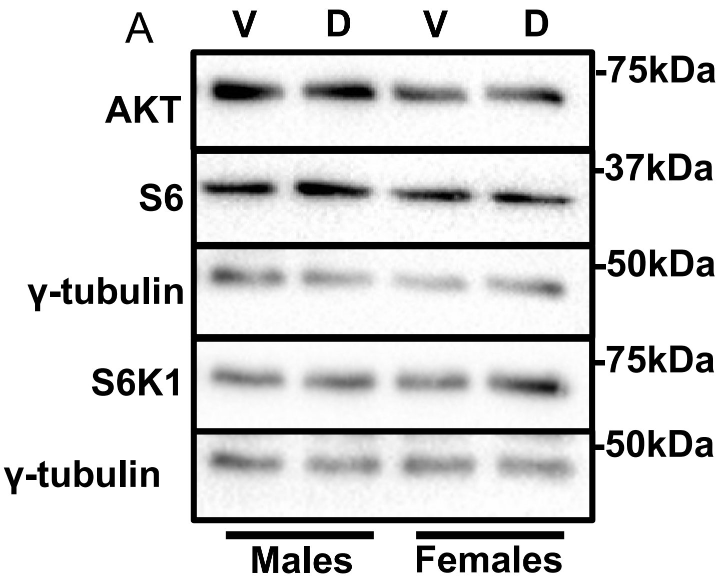


**S2 Fig, related to Manuscript Fig 2. Total protein levels of signaling proteins are not affected by chemotherapy treatment.** Male and female (18±2 months of age) CD2F1 mice were treated as described in supplementary figure 1. Immunoblots are shown for the total levels of AKT, S6 and S6K1 in the gastrocnemius **(A)**. V, vehicle; D, drug.

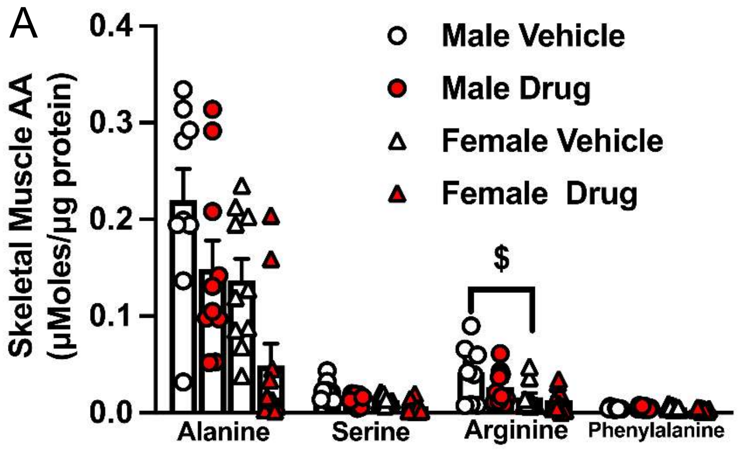


**S3 Fig. Related to Manuscript Fig 3. Minimal changes were found for other amino acids measured.** Male and female (18±2 months of age) CD2F1 mice were treated as described in supplementary figure 1. Amino acids were measured by HPLC in the gastrocnemius **(A)**. Data are mean ± SE, n = 8 – 10. Effect of drug and sex interaction: $ $p < 0.05$. Data were analyzed using a two-way ANOVA followed by a Tukey's post hoc test.

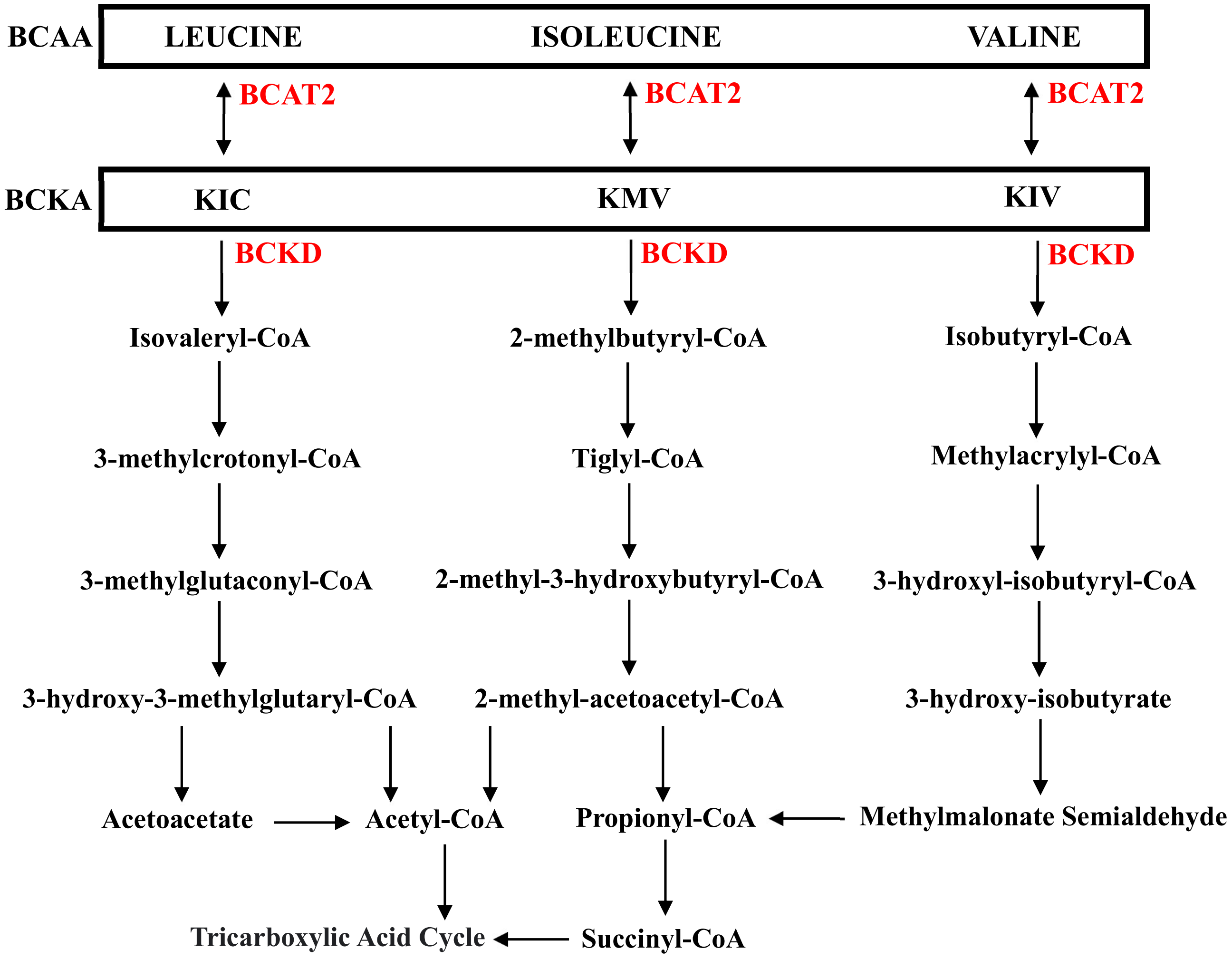


**S4 Fig Related to Manuscript Fig 4. A simplified diagram of BCAA catabolism in skeletal muscle.** The BCAA are first reversibly transaminated by BCAT2. The resulting BCKA are then irreversibly decarboxylated by the BCKD complex, yielding isovaleryl- CoA, 2-methylbutyryl-CoA and isobutyryl-CoA. Each of these is then funneled into their respective metabolic pathways. BCAA, branched-chain amino acid ; BCKA, branched-chain α-ketoacids; BCAT, branched-chain aminotransferase; BCKD, branched-chain α-keto acid dehydrogenase complex; KIC, 2-keto-isocaproate/4-methyl-2-oxopentanoic acid; KMV, α-keto-β-methylvaleric acid/3-methyl-2-oxopentanoate; KIV, 2-keto-isovalerate/3-methyl-2-oxobutanoic acid. Re-drawn and modified from Mann et al (46).

24. **Nunes EA**, **Kuczera D**, **Brito GAP**, **Bonatto SJR**, **Yamazaki RK**, **Tanhoffer RA**, **Mund RC**, **Kryczyk M**, **Fernandes LC**. β-Hydroxy-β-methylbutyrate supplementation reduces tumor

growth and tumor cell proliferation ex vivo and prevents cachexia in Walker 256 tumor-bearing rats by modifying nuclear factor-κB expression. *Nutrition Research* 28: 487–493, 2008.

34. **Englund DA**, **Jolliffe AM**, **Hanson GJ**, **Aversa Z**, **Zhang X**, **Jiang X**, **White TA**, **Zhang L**, **Monroe DG**, **Robbins PD**, **Niedernhofer LJ**, **Kamenecka TM**, **Khosla S**, **LeBrasseur NK**.

Senotherapeutic drug treatment ameliorates chemotherapy-induced cachexia. *JCI Insight* 9, 2024.
